# The 24-Hour Time Course of Integrated Molecular Responses to Resistance Exercise in Human Skeletal Muscle Implicates *MYC* as a Hypertrophic Regulator That is Sufficient for Growth

**DOI:** 10.1101/2024.03.26.586857

**Authors:** Sebastian Edman, Ronald G. Jones, Paulo R. Jannig, Rodrigo Fernandez-Gonzalo, Jessica Norrbom, Nicholas T. Thomas, Sabin Khadgi, Pieter Jan Koopmans, Francielly Morena, Calvin S. Peterson, Logan N. Scott, Nicholas P. Greene, Vandre C. Figueiredo, Christopher S. Fry, Liu Zhengye, Johanna T. Lanner, Yuan Wen, Björn Alkner, Kevin A. Murach, Ferdinand von Walden

## Abstract

Molecular control of recovery after exercise in muscle is temporally dynamic. A time course of biopsies around resistance exercise (RE) combined with -omics is necessary to better comprehend the molecular contributions of skeletal muscle adaptation in humans. Vastus lateralis biopsies before and 30 minutes, 3-, 8-, and 24-hours after acute RE were collected. A time-point matched biopsy-only group was also included. RNA-sequencing defined the transcriptome while DNA methylomics and computational approaches complemented these data. The post-RE time course revealed: 1) DNA methylome responses at 30 minutes corresponded to upregulated genes at 3 hours, 2) a burst of translation- and transcription-initiation factor-coding transcripts occurred between 3 and 8 hours, 3) global gene expression peaked at 8 hours, 4) ribosome-related genes dominated the mRNA landscape between 8 and 24 hours, 5) methylation-regulated *MYC* was a highly influential transcription factor throughout the 24-hour recovery and played a primary role in ribosome-related mRNA levels between 8 and 24 hours. The influence of MYC in human muscle adaptation was strengthened by transcriptome information from acute MYC overexpression in mouse muscle. To test whether MYC was sufficient for hypertrophy, we generated a muscle fiber-specific doxycycline inducible model of pulsatile MYC induction. Periodic 48-hour pulses of MYC over 4 weeks resulted in higher muscle mass and fiber size in the soleus of adult female mice. Collectively, we present a temporally resolved resource for understanding molecular adaptations to RE in muscle and reveal MYC as a regulator of RE-induced mRNA levels and hypertrophy.

## Introduction

Molecular alterations after a bout of exercise in skeletal muscle precede adaptation and ultimately contribute to a change in phenotype ^1–7^. Initial time course work in humans that utilized ≥3 post-exercise muscle biopsies established that the 2–4 hour recovery time point is ideal for studying targeted changes in mRNA levels after a bout of exercise ^8–11^. Others that leveraged more comprehensive temporal profiling of global gene expression ^12–14^ demonstrated that many genes have delayed and/or biphasic responses to exercise in muscle that extend well beyond 4 hours. Recent work in skeletal muscle further emphasizes that gene expression data from a single time point after exercise is limiting when trying to capture the complex and dynamic nature of the adaptive response, and could even lead to inaccurate or misleading conclusions ^15^. It is also important to consider the effects of the muscle biopsy ^11^ and circadian rhythm ^14^ in human exercise studies; these are typically unaccounted for. There is a critical need for temporally resolved and biopsy-only controlled investigations to explore and understand the molecular responses to resistance exercise since muscle mass and/or function is strongly associated with all-cause mortality ^16–19^. A detailed understanding of the most influential molecular factors during the post-resistance exercise recovery period will help focus efforts at developing targeted therapies against muscle mass loss and/or enhancing hypertrophic responsiveness to exercise.

Several seminal ^20–22^ and recent studies ^23,24^ suggest that the transcription factor *c-Myc* (referred to as *Myc* or *MYC* for mouse and human genes, respectively) is a key component of skeletal muscle hypertrophic adaptation to loading in animals. Our work using human skeletal muscle biopsies after a bout of resistance exercise (RE) ^25^, as well as meta-analytical information that combines numerous human muscle gene expression datasets during the recovery period after exercise ^26^, indicates that *MYC* is highly responsive to hypertrophic loading ^27^. MYC protein accumulates in human muscle following a bout of RE ^28–31^ as well as in response to chronic training ^32^. Its expression may also differentiate between low and high hypertrophic responders ^32^. *Myc* is induced cell-autonomously in myotubes by electrical stimulation *in vitro* ^33^ and is strongly upregulated in murine myonuclei during mechanical overload ^24^. MYC protein localizes to myonuclei during loading-induced hypertrophy ^20,21^, is considered pro-anabolic ^34–36^, and can drive muscle protein synthesis and ribosome biogenesis in skeletal muscle ^31,37–39^. Loss of MYC results in lower muscle mass in preclinical models ^40,41^. It is also estimated to target ∼15% of the genome in multiple species ^42^. Still, the magnitude of its contribution to the exercise response in humans is not well-understood ^43^, nor its sufficiency for muscle hypertrophy.

The current investigation details the global gene expression response to a bout of RE after 30 minutes, 3-, 8-, and 24-hours using RNA-sequencing (RNA-seq) in human skeletal muscle biopsy samples. We reveal the effect of the muscle biopsy and inherent circadian rhythmicity using a biopsy-only timepoint and feeding- matched control group. We then analyzed the human muscle methylome at 30 minutes after RE and combined these data with the transcriptome response to RE using a novel -omics integration approach. Integration of methylomics and transcriptomics sheds light on the molecular regulation of gene expression during recovery from exercise. With our transcriptome data, we infer the major transcriptional regulators of the exercise response using *in silico* ChIP-seq ^44^ that we have previously validated in skeletal muscle with a genetically modified mouse ^24,27^. Our molecular and computational analyses identified *MYC* as one of the most influential transcription factors on the exercise transcriptome throughout the time course of recovery from RE. Muscle-specific *Myc* overexpression data from the plantaris ^24^ and soleus ^27^ of mice was used to reinforced the human exercise data. We employed a genetically modified mouse model to induce MYC in a pulsatile fashion specifically in skeletal muscle over four weeks to determine if MYC is sufficient for hypertrophy. Our genetically driven pulsatile approach avoids potential negative effects of chronically overexpressing a hypertrophic regulator ^45,46^ and more closely mimics the transient molecular effects of exercise in skeletal muscle ^5–7,47^. This work collectively illustrates the molecular landscape with temporal resolution after a bout of RE and places *MYC* at the center of the skeletal muscle RE response in mice and humans.

## Results

### Biopsy time course at rest and the transcriptional regulation of circadian genes in human skeletal muscle

For the analysis of RNA-seq data, we focused on protein-coding genes. The number of protein-coding differentially expressed genes (DEGs, adj. *p*<0.05) during the time course of recovery (Figure 1A) in the repeated biopsies control group (CON, n=5) relative to the Pre time point (morning biopsy; Figure 1A) was: 30 minutes - 0 upregulated, 1 downregulated; 3 hours - 12 upregulated, 30 downregulated; 8 hours - 55 upregulated, 75 downregulated; and 24 hours - 0 upregulated, 1 downregulated (Figures 1B & D). The biopsy timepoints, and their naming are chosen to precisely correspond to the post RE timepoints for the RE group (Figure 2A). The Pre muscle biopsies were taken 15 minutes prior to the 45-minute control protocol (equivalent to the 45 min of resistance exercise in RE group). Thus, in practice, the biopsy named 30 minutes post is taken 90 minutes after the Pre biopsy (15 min plus, 45 min, plus 30 min), the biopsy named 3 hours is taken 4 hours after the Pre biopsy, and so on. Previous work involving human skeletal muscle biopsies revealed the regulation of circadian genes over 24 hours ^48^. We found that *NR1D1* (*REVERBα*) (adj. *p*=0.01x10^-5^), *PER1* (adj. *p*=0.0008), and *PER2* (adj. p=0.07x10^-7^) were lower at the 3-hour timepoint (Figure 1C). At 8 hours, *NR1D2* (*REVERBβ*) (adj. *p*=0.005), *PER1* (adj. p=0.03x10^-6^), *PER2* (adj. *p*=0.007x10^-10^), and *PER3* (adj. p=0.007x10^-3^) were lower (Figure 1C). *KLF15*, a circadian-regulated mediator of lipid metabolism ^48^, was lower at the 3-hour (adj. *p*=0.0001) and 8-hour timepoints (adj. p=0.03x10^-12^) in the CON group (Figure 1C). *PPARGC1β*, another circadian-controlled gene ^49^, was upregulated at 3 hours (adj. *p*=0.048) and 8 hours (adj. *p*=0.01; Figure 1C). Also worth mentioning is that *FOXO3*, a central regulator of autophagy and mass in skeletal muscle ^50–52^, was lower at 3 hours (adj. *p*=0.0004) and 8 hours (adj. *p*=0.007x10^-3^, Figure 1C). *SESN1*, which also regulates muscle mass ^53^, was lower at 3 hours (adj. p=0.003) and 8 hours (adj. p=0.008x10^-3^) (Figure 1C).

**Figure 1.**
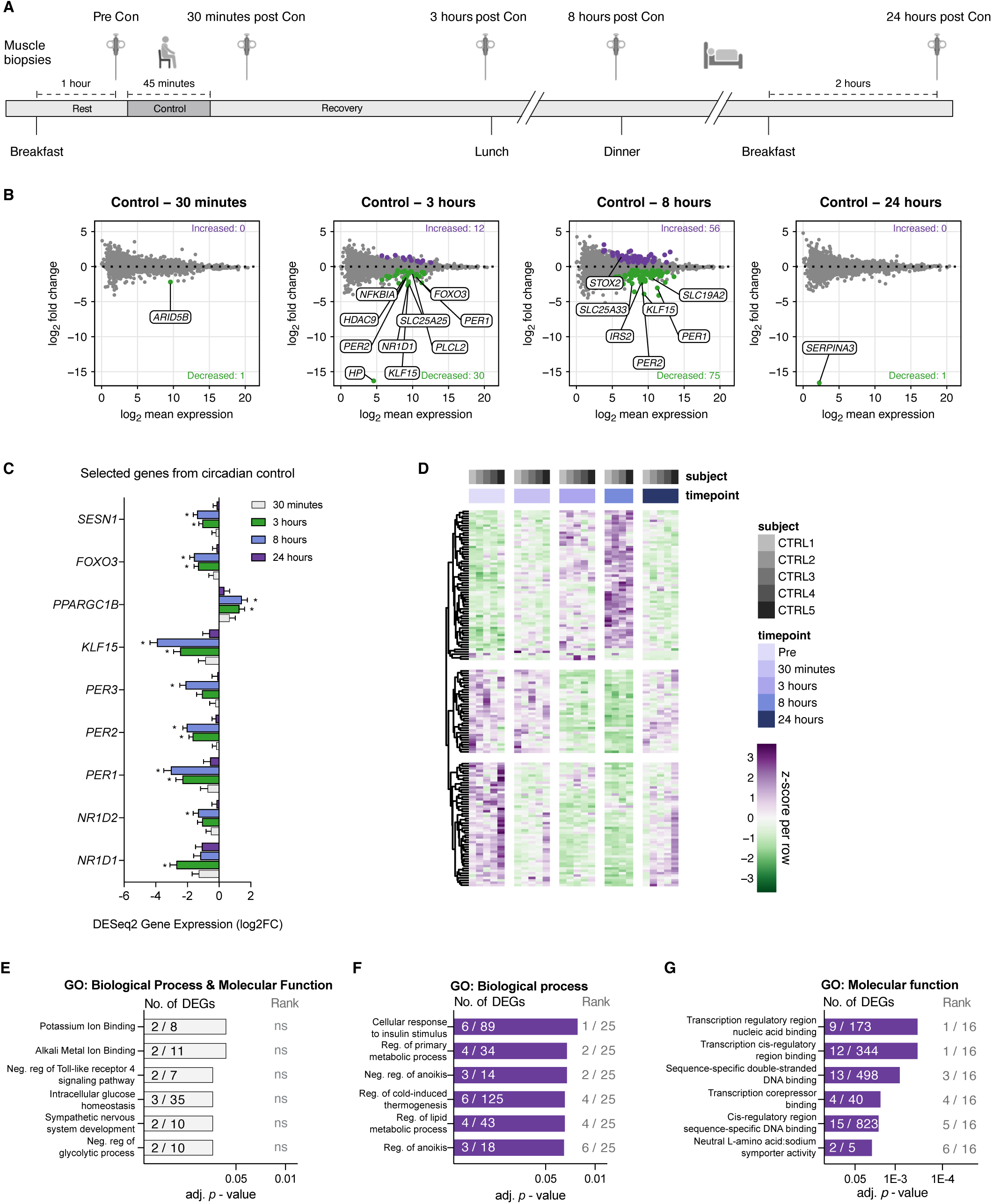
Gene expression patterns for circadian control. (A) Schematic overview for the control arm of the human intervention, n=5. (B) MA plots showing differentially expressed genes (DEG) vs pre-values, time matched to 30 minutes, 3, 8, & 24 hours of recovery in the resistance exercise trial (see Figure 2A). Purple and green dots indicate up- or down-regulated regulated genes (adj. *p*<0.05), respectively. Top genes for adj. *p*- value are highlighted in plots. (C) Fold-change for targeted DEGs in the control group. * = adj. *p*<0.05. (D) Heatmap showing z-scores for 60 up-, and 90 down-regulated DEGs across all time points and volunteers in the control trial. (E-F) Gene ontology (GO) pathway enrichment analysis on DEGs across the entire 24 h control period. Numbers within the bars indicate the proportion of DEGs in our data set corresponding to the specific pathway. Rank values indicate the specific pathways adj. *p*-value rank. E) Up-regulated Biological Processes and Molecular Functions, F) Down-regulated Biological Functions, G) Down-regulated Molecular Functions. Con = Control situation, ns= not significant, Neg. = Negative, Reg. = Regulation.

**Figure 2.**
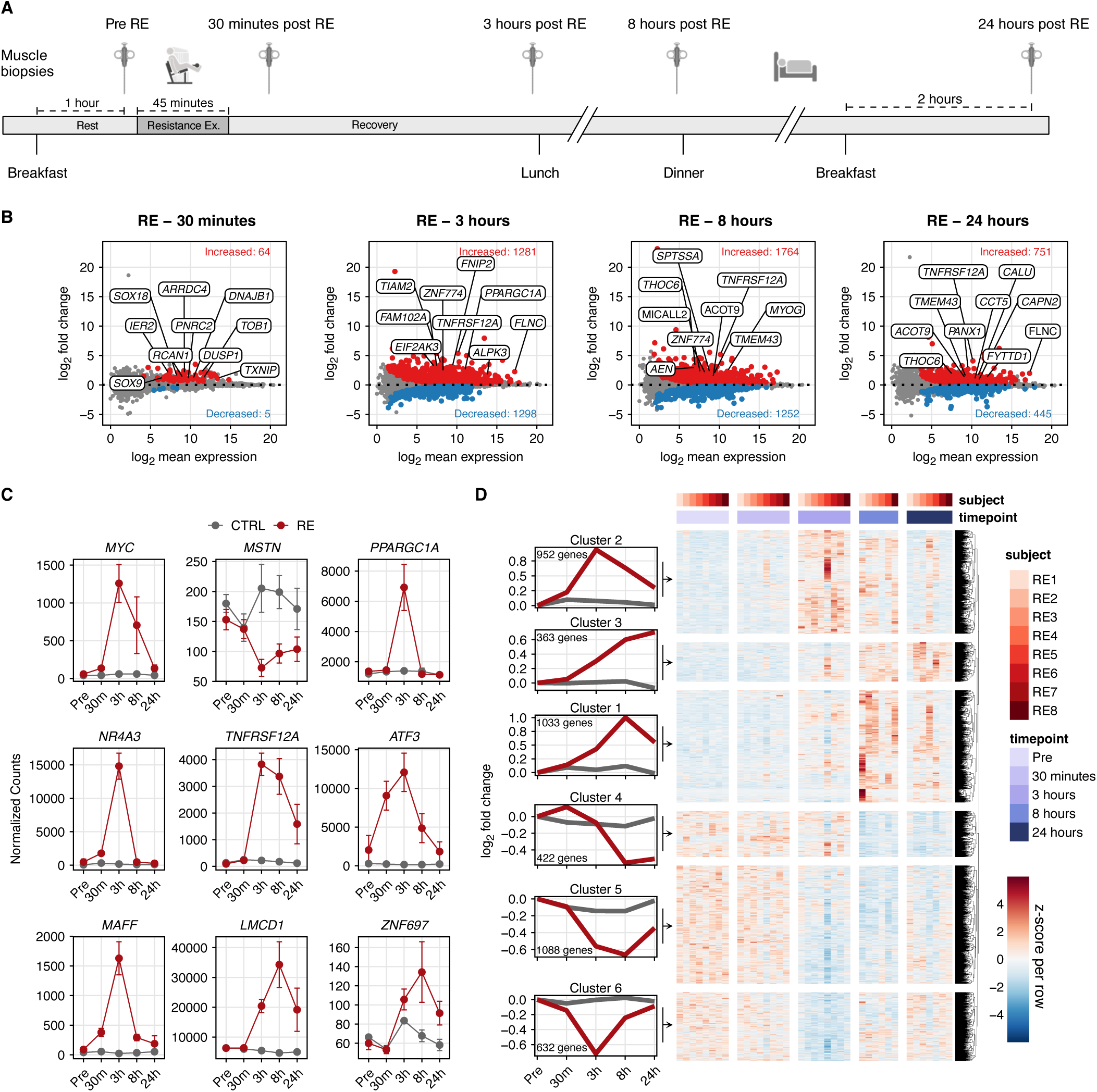
Gene expression patterns during 24 hours of recovery from resistance exercise. (A) Schematic overview of resistance exercise (RE) intervention, n=8. (B) MA plots showing differentially expressed genes (DEG) vs resting pre-values, following 30 minutes, 3, 8, & 24 hours of recovery from RE. Red and blue dots indicate up- or down-regulated regulated genes (adj. *p*<0.05), respectively. Top 10 genes for adj. *p*-value are highlighted in plots. (C) Normalized counts for targeted genes across the 24-hour intervention. Red dots = RE trial, grey dots = Control trial. (D) Heatmap showing z-scores for 2399 up-, and 2126 down-regulated DEGs across all recovery time points and volunteers in the RE trial. Genes are clustered according to their expression pattern across timepoints within the 24-hour recovery period.

To more broadly investigate what specific functions were being regulated as a consequence of our CON intervention, we ran background corrected pathway enrichment analysis using Enrichr with the 2023 gene ontology (GO) database as our cross reference (GO: biological process & molecular function) ^54–58^. Collectively, no significantly upregulated pathways were detected (Figure 1E). A few negatively regulated pathways were detected, indicating a reduced insulin stimulus and reduced transcriptional speed (Figure 1F-G). Although significantly enriched, only 2-15 genes were coding for each pathway, suggesting the effect was small.

Nevertheless, it should be noted that the significantly downregulated pathways are almost exclusively driven by DEGs expressed at 3 and 8 hours. In addition to circadian regulation, DEGs across time points in CON could be related to feeding since the Pre, 30 minutes, and 24 hours biopsies were taken 1, 2 and 2 hours after feeding, respectively, while the biopsies at 3 and 8 hours were both taken approximately 5 hours after food intake (Figure 1A). Thus, our finding of slightly downregulated insulin stimulation and transcription at 3 and 8 hours seems intuitive.

### Differentially expressed genes (DEGs) peaked 8 hours after resistance exercise (RE) relative to Pre

Following an acute bout of resistance exercise (RE, n=8, Figure 2A), the number of DEGs relative to the Pre time point (adj. p<0.05) was: 30 minutes - 64 upregulated, 5 downregulated; 3 hours - 1281 upregulated, 1298 downregulated; 8 hours - 1764 upregulated, 1253 downregulated; and 24 hours - 751 upregulated, 445 downregulated (Figure 2B). As anticipated, we noted substantial expression changes in genes previously recognized as responsive to RE and/or important for muscle remodeling (Figure 2C) ^25,26,59,60^. Of all upregulated genes across the 24-hour recovery period in the RE group, 46% were differentially expressed at two or more time points while the proportion was 34% for downregulated genes. In total, 2399 upregulated and 2126 downregulated unique DEGs were identified throughout the 24-hour recovery period (Figure 2D).

### The integrated 24-hour transcriptome time course of recovery after acute RE

Using the two lists of all DEGs from across the entire 24-hour recovery period after RE (up- or down-regulated) relative to Pre generated in Figure 2D, we ran background corrected pathway analysis as described above ^54–58^. For the 2399 up-regulated DEGs, 44 biological processes (adj. *p*<0.05) were identified, with a large portion of the pathways in the 24-hour post-RE window relating to transcription, translation, and the synthesis of new ribosomes. After exclusion of pathways with large overlaps in underlying DEGs, the top (adj. *p*-value ranked) processes were ribosome biogenesis (GO:0042254), activation of protein localization to telomere (GO:1904816), inhibition of apoptosis (GO:0043066), activation of transcription by RNA polymerase II (GO:0045944), activation of intracellular signal transduction (GO:1902533), and response to unfolded protein (GO:00066986) (Figure 3A). Fourteen molecular function pathways were also identified (adj. *p*<0.05). Of the molecular functions identified, RNA binding (GO:0003723), cadherin binding (GO:0045296), ubiquitin protein ligase binding (GO:0031625), purine ribonucleoside triphosphate binding (GO:0035639), protein phosphatase 2A binding (GO:0051721), and MAP kinase tyrosine/serine/threonine phosphatase activity (GO:0033550) were the top pathways, again excluding pathways with large overlap (Figure 3B).

**Figure 3.**
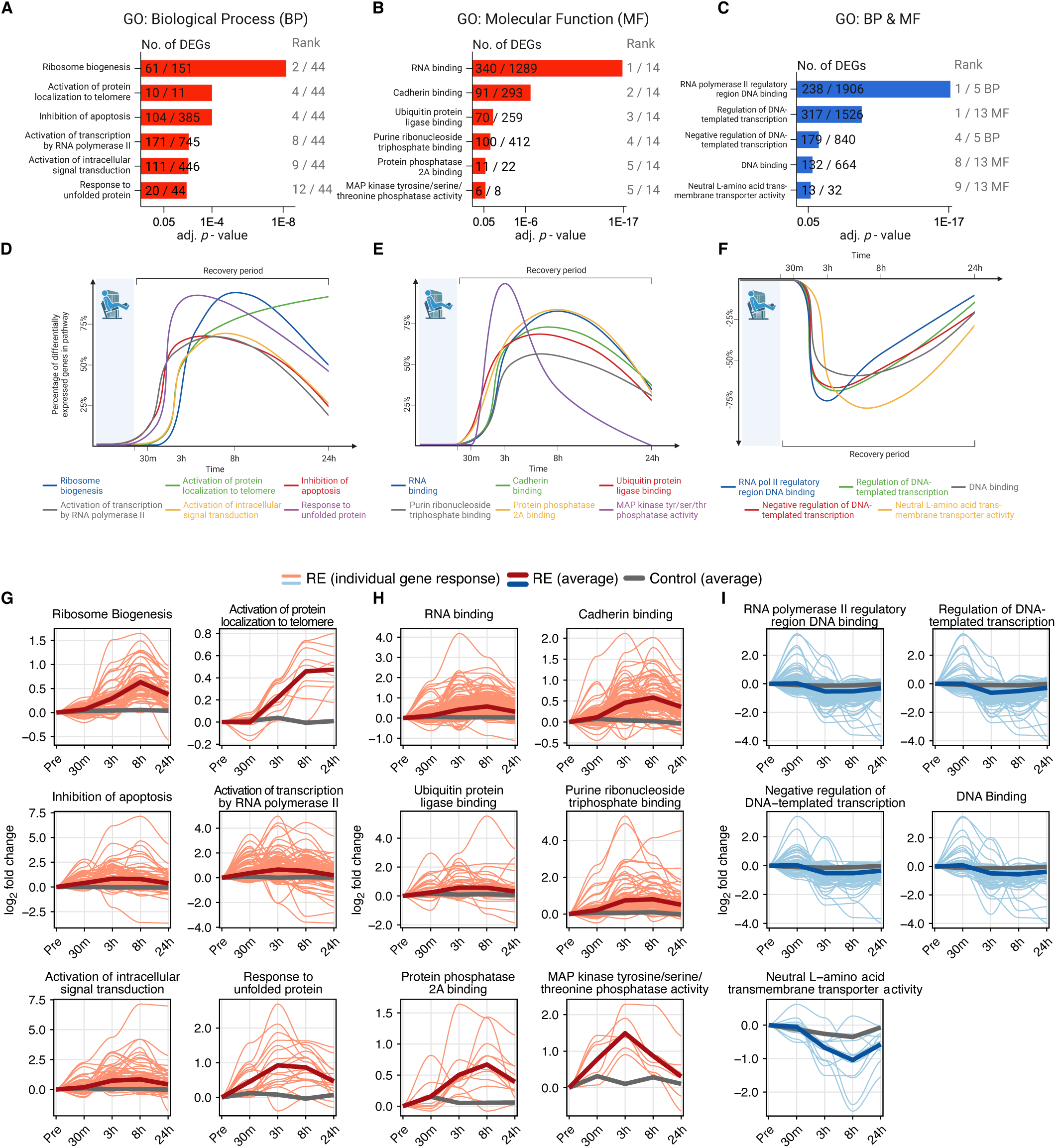
Pathway enrichment time course across 24 hours of recovery from resistance exercise. (A-C) Gene ontology (GO) pathway enrichment analysis on DEGs across the entire 24-hour recovery period. Numbers within the bars indicate the proportion of DEGs in our data set corresponding to the specific pathway. Rank values indicate the specific pathways adj. *p*-value rank. The corresponding timeline shows the proportion of DEGs with the specific pathways across the 24-hour recovery period. A) Upregulated Biological Processes, B) Up-regulated Molecular function, and C) Down-regulated Biological Processes and Molecular function. (D-F) Timelines of the top (GO) pathways from A-C expressed as a percentage of the number of DEGs within that pathway. D) Upregulated Biological Processes, E) Up-regulated Molecular function, and F) Down-regulated Biological Processes and Molecular function. (G-I) Average fold-change (vs Pre) for highlighted pathways (thick red lines) along individual DEGs within the specific pathway (lighter red lines). The average fold-change for the same genes in the control situation is presented in grey. G) Up-regulated Biological Processes, H) Up- regulated Molecular function, I) Down-regulated Biological Processes and Molecular Function.

Next, we identified at which timepoint all DEGs within each specific pathway were differentially regulated relative to Pre. The number of DEGs at each timepoint was then expressed as a percentage of the entire pathway response (e.g. 61 DEGs in our dataset were found to regulate ribosome biogenesis, and of these 61 genes, 57 – or 93% - were expressed following 8 hours of recovery). Plotting these values for all time points relative to Pre thus reveals a 24-hour temporal pattern of each pathway following acute RE (Figure 3D & E). None of the most highly enriched pathways within our analysis peaked at 30 minutes post-exercise. However, following a targeted analysis of enriched pathways with peak expression at 30 minutes, we observe that growth factor- and glucocorticoid responses are strongest at 30 minutes post-RE, as well as stress response signaling and mRNA catabolism (Supplemental Figure 1A). The peak in transcripts coding for mRNA catabolism 30 minutes after RE (*BTG2*; adj. *p*=0.005, *ZC3H12A*; adj. *p*= 0.003, *ZFP36L1*; adj. *p*= 0.01, and *TOB1*; adj. *p*=0.0005) precedes any marked downregulation of DEGs, suggesting major catabolism of mRNA occurs in muscle following upregulation of anti-proliferative- and mRNA-decaying enzymes.

At 3 hours post RE relative to Pre, we observed two major upregulated pathways peaking: response to unfolded proteins (20/44 genes; Figure 3A & D) and MAP kinase phosphatase activity (6/8 genes; Figure 3B & E). The former of the two is primarily driven by genes coding the heat shock protein family, such as *DNAJA1* (adj. *p*= 8.5x10^-7^ at 3 hours) and *HSPA1A* (adj. *p*= 8.5x10^-5^ at 3 hours). The latter, MAP kinase phosphatase activity, is driven by genes encoding the dual specificity phosphatase protein family (DUSP), responsible for dephosphorylation of tyrosine/serine/threonine sites (*DUSP2, 4, 5, 8, 10* & *16*, adj. *p*=0.05 at 3 hours). DEGs within this pathway peak at the 3-hour timepoint, with only two remaining elevated at 8 hours. This pattern was also reflected when mapping the fold change of DEGs within each pathway, rather than the number of DEGs, across the 24-hour recovery (Figure 3G-H). A clear peak in genes encoding phosphatase activity directed towards the MAP kinase superfamily early during RE recovery may be a response triggered by the rapid severalfold increase in protein phosphorylation of mTOR-targets such as S6K1 and 4EBP1 occurring at around 60-90 minutes post RE ^61,62^. According to this previous work, the rapid rise in anabolic signaling via protein phosphorylation at this time point is then followed by a swift decrease, with some signaling proteins showing close to baseline phosphorylation levels at 3 hours of recovery.

Several upregulated pathways are overrepresented to a similar degree at 3 and 8 hours of recovery such as ubiquitin protein ligase binding (70/259 genes; GO:0031625; Figure 3B, E & H), activation of transcription by RNA polymerase II (171/745 genes; GO:0045944; Figure 3A, D & G), and activation of intracellular signal transduction (111/446 genes; GO:1902533; Figure 3A, D & G), pointing to increased protein turnover. At the same time, inhibition of apoptosis (104/385 genes; GO: 0043066; Figure 3A, D & G) was also upregulated, which may be a direct response to the increased transcriptional emphasis on the ubiquitin system. The post-translational modifications mediated by ubiquitination of pro-apoptotic Bcl-2 and BH3-only proteins have been proposed to be crucial for cell survival ^63^. For instance, the transcript *RNF144B* coding for the E3 ubiquitin ligase *IBRDC2*, which targets the Bcl-2 ‘executioner’ Bax for ubiquitination ^64^, is significantly upregulated at 3 hours only (adj. p=2.8x10^-7^).

Following the burst of transcription- and translation initiation-coding transcripts at 3 and 8 hours, the mRNA landscape shifted toward the ribosome. At 8 and 24 hours of recovery, regulation of ribosome biogenesis (61/151 genes; GO:0042254; Figure 3A, D & G) and RNA binding (340/1289 genes; GO:0003723; Figure 3B, E & H) appear to be the dominant pathways. Within the RNA binding pathway, genes supporting ribosome assembly, posttranslational control of RNA, splicing via RNA-binding protein family (e.g. *RBMX*, *RBM15*, *RBM39*), heterogeneous nuclear ribonucleoproteins (e.g. *HNRNPU*, *HNRNPR*, *HNRNPC*) and zinc fingers (e.g. *ZNF326*, *ZNF579, ZNF697*) were differentially expressed. Moreover, several transcripts coding ribosome biogenesis factors (e.g. *BMS1* and *LTV1*) as well as ribosomal assembly and transport proteins (e.g. *NIP7*, *NOP14*, *RPF2*) composed the highly enriched ribosome biogenesis pathway (GO:0042254).

Pathways enriched within the 2126 down-regulated DEGs were considerably fewer compared to the up- regulated genes (Figure 3C, F & I). Here, five biological processes and 13 molecular functions were down- regulated (adj. *p*<0.05). Out of the five significant biological processes, four were related to transcription. The majority of DEGs within these transcription-related pathways were classified as inhibitors of transcription, meaning RE likely acts on transcription by up-regulating activation (Figure 3A) and by repressing its suppressor genes (Figure 3C) to a similar extent. In addition to transcription, histone H3 methyltransferase activity was one of the pathways found to be significantly downregulated at 3 and 8 hours following RE using a targeted analysis of these specific time points (Supplemental Figure 1B).

Specifically focusing on genes that were upregulated early after RE and downregulated later relative to Pre (Cluster 4; Figure 2D), we found that some of the negative regulators of RNA Pol II transcription (GO:0045892, GO:0000122) followed this pattern - upregulated 30 minutes and/or 3 hours while downregulated later at 8 and 24 hours (Supplemental Figure 1C). Nuclear receptor subfamily 4 group A genes *NR4A1* and *NR4A2* were among the 10 transcripts in the sequence-specific DNA binding pathway (GO: 0043565) that most clearly followed a biphasic expression pattern (up early, down late; Supplemental Figure 1D). Related to *NR4A1* and *NR4A2*, *NR4A3* showed similar biphasic tendencies, being upregulated at 3 hours (4.93 logFC, adj. p<0.05) and numerically downregulated at 24 hours relative to Pre (-0.78 logFC, adj. *p*>0.05; Supplemental Figure 1D). This finding is in agreement with previous reports suggesting *NR4A* family transcripts are highly responsive early during exercise recovery ^26,65^. Others have also shown that *NR4A3* is upregulated by seemingly opposing stimuli – following both acute exercise as well as long term inactivity ^26,65^. As many acute exercise interventions only sample muscle for 3-5 hours during recovery, a delayed depression of the *NR4A3* gene (at 24 hours into recovery) due to the biphasic nature of the *NR4A* gene family (Supplemental Figure 1D) may be underappreciated. An interpretation could be that exercise ubiquitously drives *NR4A* expression, while in fact, the post-exercise induction and repression could be balanced, and this pattern may have a specific biological function pertaining to exercise adaptation. Regardless, the existence of biphasic genes within 24 hours of RE recovery highlights the importance of considering muscle biopsy sampling time points when interpreting data.

### Changes to the muscle DNA methylome at 30 minutes of recovery after RE associates with mRNA responses at 3 hours

Binding and expression target analysis (BETA) is a multi-omics integration method for understanding transcriptional regulation ^66^. We recently adapted this method for combining reduced representation bisulfite sequencing (RRBS) data with RNA-sequencing data to understand how DNA methylation regulates the transcriptome during an acute loading stimulus in mice ^67^. Briefly, BETA considers relative methylation status (both hypo- and hyper-methylation) in relation to transcription start sites using weighted scores to infer transcriptional regulation, which is then combined with transcriptomic data for validation. In our recent work, myonuclear DNA methylation status coincided with changes in myonuclear gene expression as well as the acute metabolic responses that occurred during rapid muscle growth, giving us confidence in the validity of BETA ^67^. We leveraged RRBS and RNA-sequencing data in the current study to provide a deeper understanding of transcriptional regulation in response to acute RE.

We first used BETA to compare the methylome and transcriptome responses to RE at 30 minutes of recovery versus Pre. Combining datasets at this time point revealed <10 genes were likely being regulated at the level of methylation. This result seems intuitive since changes in DNA methylation typically precede changes in gene expression ^68^. As such, we combined the 30 minutes methylome data with the transcriptome response to RE. Changes to the methylome 30 minutes after RE were strongly predictive of the changes observed in gene expression at 3 hours after RE versus Pre (Figure 4A). This analysis inferred significant methylation control of 936 upregulated genes (*p*=0.000007). Of upregulated genes with a coordinated methylome and transcriptome response, *TNFRSF12A* (*FN14*) was the most significant (p=0.000035; Figure 4B). Upregulation of the TWEAK receptor *Fn14* occurs during the muscle hypertrophic response to exercise specifically in fast-twitch type 2 fibers of humans ^69,70^. This role for *FN14* induction during muscle adaptation could be related to non-canonical NF-κB pathway activation ^71^. Furthermore, inhibition of *Fn14* in human myotubes increases C/EPβ and MuRF ^72^. Alternatively, mechanistic work in rodents suggests *Fn14* knockout in muscle fibers improves endurance exercise capacity and inhibits neurogenic muscle atrophy ^73^, but ablation in satellite cells attenuates muscle regeneration ^74^. More gain and loss of function studies are needed to clarify the role of *Fn14* in hypertrophic muscle adaptation ^75,76^. Other notable genes with a coordinated upregulated response to RE include: *RUNX1* (Figure 4B), which regulates muscle mass ^77^ and is enriched in myonuclei during rapid load-induced hypertrophy ^24^; *RBM10* (Figure 4B), an RNA splicing factor that we previously showed is altered at the methylation level in muscle with late-life hypertrophic exercise in mice ^76,78^; and *NR4A3* (Figure 4B), among the most exercise-responsive genes in skeletal muscle that controls metabolism ^26^. We previously reported that promoter region DNA hypomethylation of *Myc* in myonuclei ^79^ coincided with strong upregulation of myonuclear and tissue *Myc* levels during acute mechanical overload in mice ^24,79^. BETA also suggested coordinated methylation and transcriptional regulation of *MYC* by RE in human muscle here (Figure 4B). Evidence in cancer cells indeed suggests that *MYC* transcription is regulated by DNA methylation status ^80–84^, in addition to regulation by other epigenetic layers ^85,86^ and G-quadruplexes ^87^. Of the 936 methylation-controlled upregulated genes predicted by BETA, 155 were coding for five biological processes as suggested by pathway enrichment analysis (Figure 4C).

**Figure 4.**
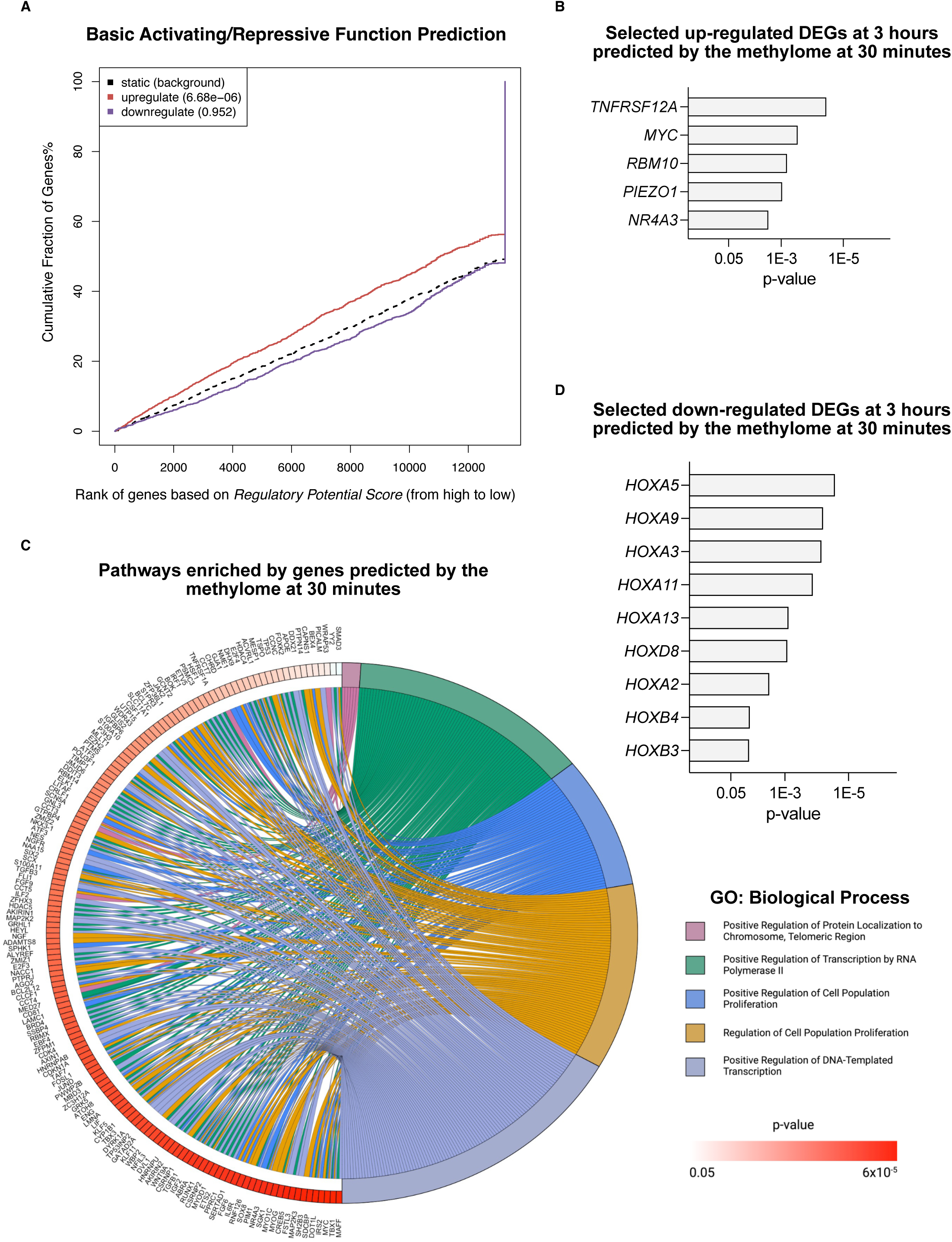
Immediate DNA-methylome changes predict transcriptional regulation. (A) Binding and expression target analysis (BETA) combining the DNA methylome at 30 minutes to differentially expressed genes (DEGs) at 3 hours post resistance exercise (RE). BETA integration analysis of up- and downregulated genes to RRBS methylation performed relative to background with predictive interaction significance represented by p-values in parenthesis. (B) Selection of up-regulated differentially expressed genes (DEGs) at 3 hours post-RE significantly predicted by the methylome at 30 minutes. (C) Chord plot illustrating the top five biological processes (gene ontology) regulated at 3 hours post exercise by genes predicted by the methylome at 30 minutes post exercise. Pathway-associated genes are ordered according to their p-values. (D) Selection of downregulated DEGs at 3 hours post RE suggested to be affected by methylation changes at 30 minutes post RE.

BETA integration of 30-minute methylome responses with 3-hour transcriptome responses to RE was not significant overall for downregulated genes. However, gene-by-gene analysis revealed methylation control of widespread downregulation of HOX genes - *HOXA2*, *HOXA3*, *HOXA5*, *HOXA9*, *HOXA11*, *HOXA13*, *HOXB3*, *HOXB4*, and *HOXD8* (Figure 4D). In muscle, HOX genes are highly regulated by DNA methylation ^88^, which serves as methylation hotspots during aging, and are affected at the methylation and mRNA levels by physical activity in humans ^89,90^. We previously reported methylation changes around HOX genes in myonuclei during hypertrophy ^91^ and with exercise during aging in muscle ^78^. HOX genes control muscle development ^92,93^, but little is known about their role in RE adaptation in adult skeletal muscle.

### MYC governs the late stage RE response via several processes

Due to the overall dominance of upregulated genes coding for ribosomal biogenesis and RNA-binding (Figure 3A & B), we asked which transcription factors may be steering transcription toward these pathways. To answer this, we first ran an epigenetic Landscape In Silico deletion Analysis (LISA) ^44^ on all upregulated genes across the 24 hour time course. We previously validated the accuracy of this computational approach for *Myc* in skeletal muscle ^24,27^. The five top transcription factors influencing the totality of the 24-hour recovery period were *NEFLA*, *BCL3*, *FOS*, *MYC*, and *ATF3*, respectively (Figure 5A). Since ribosome-related gene expression primarily dominated the late-stage recovery at 8 and 24 hours (Figure 3D, E, G & H), we modeled which transcription factors were controlling transcription for DEGs upregulated at the later time points of recovery.

**Figure 5.**
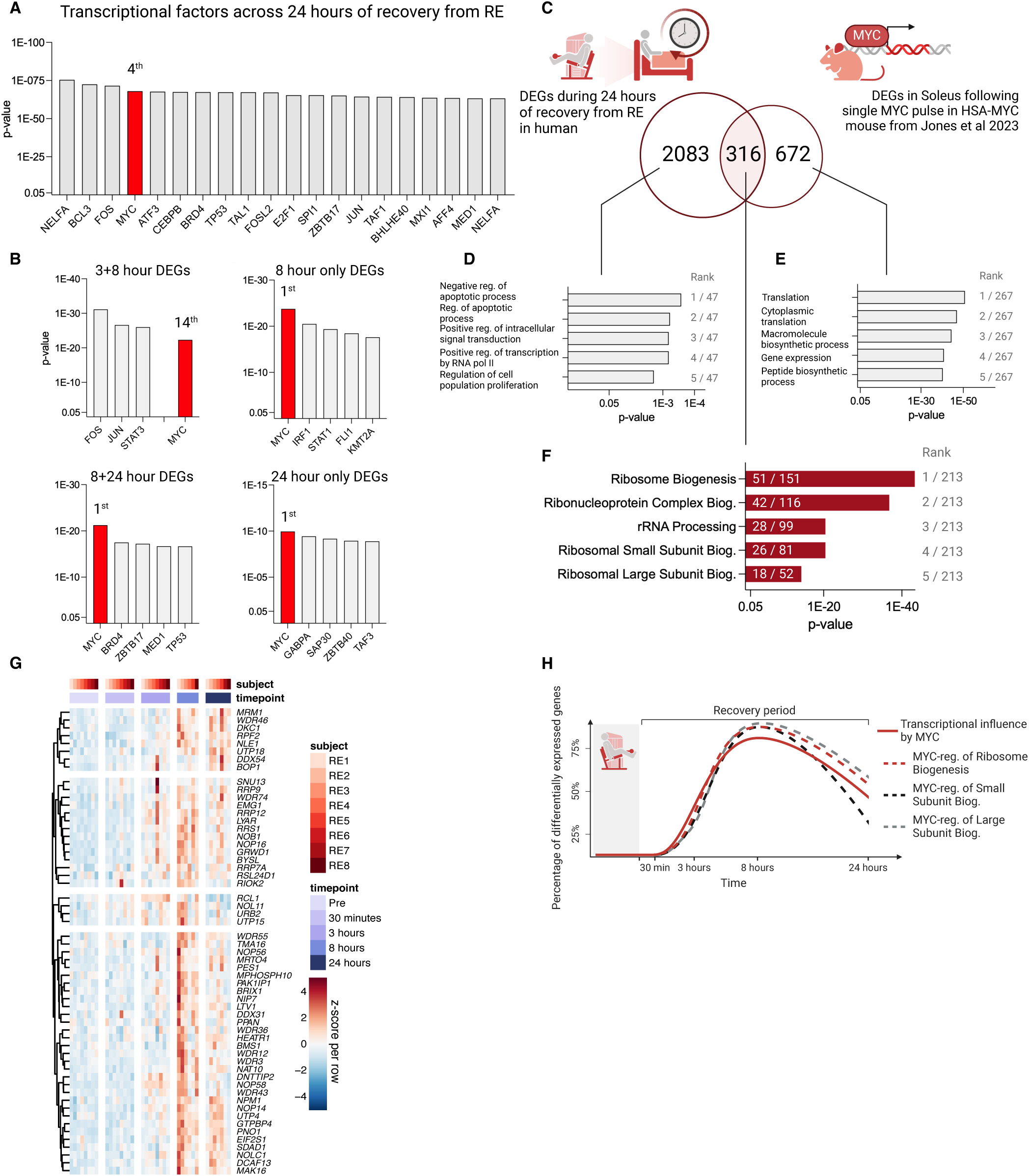
Transcription factor MYC dominates late-stage acute recovery from RE by regulating ribosome biogenesis. (A) Transcription factors predicted to be active during the 24-hour recovery period from resistance exercise (RE) sorted by p-value. (B) Transcription factors predicted to regulate the genes expressed exclusively at the later stages of acute recovery. (C) Comparison of up-regulated DEGs across 24 hours of RE recovery in humans (n=8) vs soleus muscle from MYC-overexpressing mice from Jones et al (2023). (D-F) Top five pathways (GO: Biological Processes) based on DEGs in D) the human exclusive gene list, E) MYC mouse exclusive gene list, and F) overlapping gene list, respectively. Pathways are ranked according to their adj. *p*- values. (G) Heatmap showing DEG pattern for ribosome-related genes overlapping human RE response to a MYC response in mouse soleus muscle. Genes retrieved from all five pathways presented in 3F. (H) Time courses for MYC’s transcriptional influence (Red solid line), as well as three MYC-regulated gene pathways (dashed lines).

The influence of MYC on transcription coincided with the transcriptome shift towards the ribosome (Figure 3A, B & Figure 5B, H). *MYC* was the number one transcription factor for genes expressed in the later stages of recovery - genes exclusively expressed at 8 and 24 hours relative to Pre (Figure 5B).

Next, we compared the human 24-hour post-exercise transcriptional landscape to our previously published data sets on MYC-overexpressing mice ^24,27^. Briefly, for these experiments, we generated a doxycycline-inducible muscle-specific model of pulsed MYC overexpression, called HSA-MYC (human skeletal actin reverse tetracycline transactivator tetracycline response element “tet-on” MYC) ^24,27^. Twelve hours of doxycycline in drinking water, followed by 12 hours of non-supplemented water, causes upregulation of MYC protein in skeletal muscle ^27^. MYC returns to baseline levels after 24 hours of drinking un-supplemented water (Supplemental Figure 2). Thus, we profiled the transcriptome in the plantaris and soleus muscles 12 hours after doxycycline administration ^24,27^. Of the 2399 up-regulated DEGs induced by RE over 24 hours, 316 upregulated genes overlapped the response elicited in the mouse soleus muscle by a single MYC pulse (Figure 5; Jones et al.^27^). Removing the overlapping genes from the human RE response subsequently steered the transcriptional landscape away from the ribosome, as indicated by pathway enrichment analysis (GO: biological processes) on the remaining 2083 DEGs (Figure 5D). Consequently, using the same pathway enrichment analysis on the 316 genes overlapping the human RE response and soleus transcriptome from the MYC overexpression data generated pathways largely related to the ribosome (Figure 5F). By contrast, the 672 DEGs exclusive to the MYC induction mouse mainly regulated pathways associated with acute changes to transcriptional and translational speed (Figure 5E). Regulation of similar genes was also evident, albeit to a lesser extent, when overlapping the human RE response to the smaller MYC-mediated transcriptional response in the plantaris muscle (Supplemental Figure 3; Murach et al. 2022 ^24^).

Beyond regulation of the ribosome, other genes up-regulated by both MYC and RE included those involved in actin folding by CCT/TriC (*CCT3*, *CCT7*, *CCT4*, *CCT6A*, *CCT8*, *CCT5*, *CCT2*, *TCP1*), a chaperonin complex that controls sarcomere assembly and organization in striated muscle ^94,95^. Metabolism of nucleotides (*AMPD2*, *AK6*, *GART*, *IMPDH1*, *IMPDH2*, *NME1*, *NME2*, *PPAT*, *UCK2*) and autophagy genes (*ATG3*, *HSF1*, *HSPA8*, *HSP90AA1*, *PGAM5*, *TOMM5*, *TOMM22*, *TOMM40*) were also up-regulated. Down-regulated genes shared by RE and MYC induction included *DNMT3A*, a regulator of DNA methylation in skeletal muscle^96,97^, and genes involved in ErbB signaling (*CAMK2G*, *CDKN1B*, *ERBB3*, *GAB1*). Taken together, these data suggest that MYC induction induced by acute RE in untrained humans may influence the muscle transcriptome in part by directing transcriptional machinery toward the formation of new ribosomes and/or ribosome specialization via variation in ribosomal proteins. The latter is an emerging but under-studied area of muscle hypertrophic regulation that deserves further consideration ^98^. MYC may also regulate several other processes involved in skeletal muscle exercise adaptation including proper actin folding, autophagy, and DNA methylation.

### Pulsed muscle fiber-specific MYC induction in mice is sufficient for soleus muscle hypertrophy

Our data so far suggests that MYC is a major transcriptional regulator during the acute recovery from RE in human skeletal muscle. An association between skeletal muscle hypertrophy and MYC-controlled exercise responses such as enhanced ribosome biogenesis is well-established ^25,32,39,99,100^, and inhibiting MYC in myotubes blunts ribosome biogenesis and protein synthesis ^39^. Still, it is unclear whether repeated MYC stimuli alone are sufficient to induce hypertrophy. To address this, we utilized our murine doxycycline-inducible muscle-specific model of pulsatile MYC overexpression: HSA-MYC ^24,27^.

We provided doxycycline-supplemented drinking water to 4-month-old female HSA-MYC mice for 48 hours, followed by five days of un-supplemented water for four weeks (five total MYC treatments in n=9 animals). Doxycycline-treated littermate HSA mice were controls (n=7 animals) (Figure 6A). The doxycycline treatment strategy is similar to the approach from the Belmonte laboratory for overexpressing Yamanaka factors specifically in muscle fibers ^101^. Furthermore, ∼48 hours of MYC induction in muscle approximates MYC induction in response to several exercise sessions per week.

**Figure 6.**
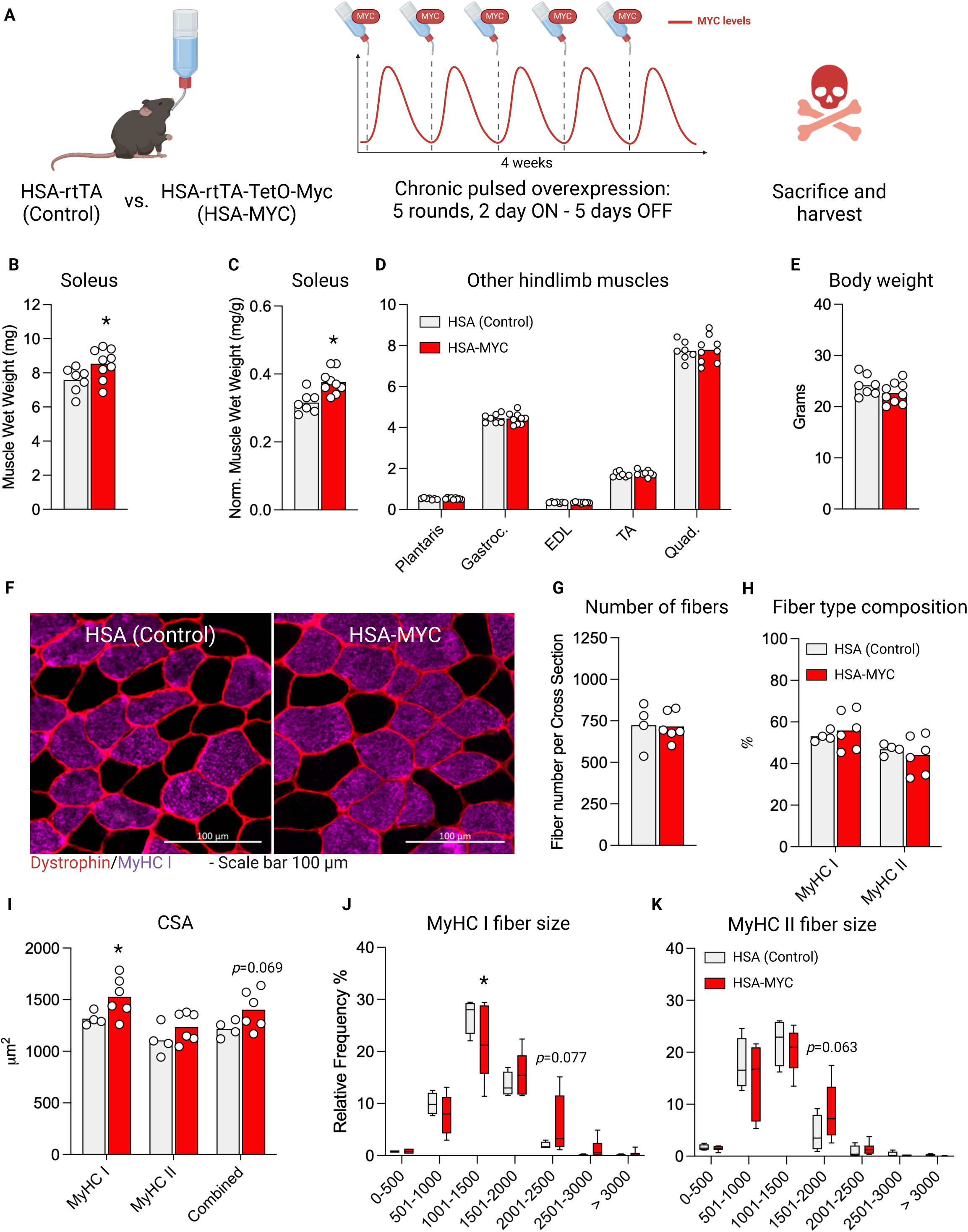
Four weeks of pulsed MYC overexpression is sufficient to cause muscle fiber type specific hypertrophy. (A) Graphical representation of the experimental design, (B) Soleus muscle weight of HSA Control and HSA MYC mice after 5 bolus exposures over 4 weeks (C) Soleus muscle wet weight normalized to body weight. (D) Hindlimb muscle weight of plantaris, gastrocnemius (gastroc.), extensor digitorum longus (EDL), tibialis anterior (TA) and quadriceps (Quad.) normalized to body weight. (E) Bodyweight of mice (F) Representative images of MyHC I (purple) and MyHC II (black) muscle fiber size in HSA Control (left) and HAS-MYC (right) mice. Dystrophin is outlining fiber borders (red). Scale bar is 100 μm. (G) Total number of fibers per soleus muscle. (H) Distribution of MyHC I and MyHC II fibers expressed as a percentage. (I) Cross sectional area MyHC I and MyHC II fibers, and their average. (J-K) Frequency distribution plot for fiber type size of MyHC I (J) and MyHC II (K) fibers. MyHC= Myosin Heavy Chain, CSA= Cross sectional area. (B-E) N= 7 HSA Control, 9 HSA MYC. (F-K) N= 4 HSA Control, 6 HSA-MYC. White dots are individual values. * = *p*<0.05.

Pulsed MYC induction resulted in a larger absolute mass (+12.5%, *p*=0.002; Figure 6B) and normalized mass (+20.7%, *p*=0.025; Figure 6C) of the soleus muscle relative to controls. The murine soleus muscle contains a myosin heavy chain (MyHC) fiber type distribution similar to the human vastus lateralis muscle (∼50% MyHC I and ∼50% MyHC IIa) ^27,102,103^, which is the muscle from which biopsies were obtained for the current study. The weight of other predominantly fast-twitch (2A, 2B, and 2X predominant) mouse hindlimb muscles was not different with MYC induction versus controls (p>0.05) (Figure 6D). Likewise, the body weight of the mice was not different between groups (p=0.49, Figure 6E) nor was food intake in a subset of mice (data not shown). These data collectively suggest a muscle and/or fiber-type dependent effect of MYC for inducing muscle hypertrophy. To further interrogate this growth, we performed immunohistochemistry on soleus muscle (Figure 6F). We observed no changes in the total amount of fibers within the soleus (Figure 6G), nor did we detect major shifts in muscle fiber type distribution (Figure 6H). However, overall (+15.1%, *p*=0.069) and MyHC I fiber cross sectional area (+16.1%, *p*=0.043) was larger with pulsatile MYC induction relative to controls (Figure 6I). A rightward shift in fiber size was also observed (Figure 6J). Fibers expressing MyHC II had a lesser response to pulsatile MYC induction, showing +11.6% difference (p=0.22; Figure 6I & K). Our prior work shows that the global transcriptional response to a single pulse of MYC is most pronounced in the soleus (∼1400 DEGs) relative to the plantaris (∼500 DEGs) and the quadriceps (<50 DEGs) ^24,27^, which likely contributes to soleus- specific mass gains. Future investigations will delve deeper into MYC dynamics across muscles and the specific mechanisms by which MYC mediates growth of the soleus. Nevertheless, we provide the first evidence that MYC is sufficient for muscle hypertrophy in the myosin fiber types expressed in human skeletal muscle.

## Discussion

The 24-hour time course of molecular responses to RE in human skeletal muscle revealed several fundamental aspects of exercise adaptation: 1) the DNA methylome response to RE at 30 minutes strongly influences global gene expression at 3 hours, 2) a burst of translation and transcription initiation coding transcripts occurs between 3 and 8 hours, 3) global gene expression peaks at 8 hours after an RE bout, 4) ribosomal-related gene expression dominates the mRNA landscape between 8 and 24 hours during recovery after RE, 5) MYC is predicted as a highly influential transcription factor throughout the 24 hour recovery period and plays a primary role in ribosomal-related transcription between 8 and 24 hours, and 6) periodic pulses of MYC are sufficient to drive muscle growth in the mixed fiber type soleus muscle of female mice.

Two to four hours after RE is typically considered the ideal time to detect changes in gene expression in skeletal muscle ^8,9,70,104,105^. By contrast, our data show that the largest number of DEGs is detected 8 hours into recovery. These new findings may influence the design of future RE studies that aim to evaluate gene expression and inform when single time-point biopsies should be taken to interrogate specific recovery processes (e.g. ribosome biogenesis versus MAPK gene expression versus ubiquitin and apoptotic gene expression). Using the same human muscle samples from this study, we previously reported that ribosome biogenesis peaks at 3 hours after RE, recovers at 8 hours, then rises again at 24 hours ^25^. The induction of ribosomal-related mRNAs between 8 and 24 hours likely relates to the biphasic increase in rRNA that may support translational capacity for muscle growth^38,106^. To this point, our data reveal unique and sometimes multimodal or divergent patterns of gene expression across gene categories over the 24-hour time course of recovery from RE. These patterns lend perspective to instances where opposite results in specific gene responses at different post-exercise time points are reported.

Previous studies report mixed findings regarding the agreement between methylation and gene expression in skeletal muscle with exercise training in humans; the relationship may be weak ^107^ or relatively strong ^108,109^. With acute exercise, however, it is shown that the acute methylation response may predict mRNA levels when specifically evaluating the promoter of exercise-responsive genes ^68,110^. Using a time course approach with high temporal resolution and a novel and holistic -omic integration technique, we show a strong relationship between the 30-minute post-exercise global methylome response and the 3-hour post-exercise transcriptome of upregulated genes in skeletal muscle from recreationally trained individuals. This relationship may change with more structured training, however. For instance, the *Myc* response to repeated bouts of RE tends to become blunted over time ^23^; this may contribute to hypertrophic response heterogeneity between individuals ^32,43,111,112^. BETA analysis predicted MYC transcription to be regulated at the epigenetic level, consistent with work in cancer cells ^80–84^. Our current and previous findings suggest that *Myc* is regulated by DNA methylation status in muscle fibers during hypertrophy^79^. More work is needed to determine whether epigenetic changes underpin lower transcriptional sensitivity of *Myc* to repeated bouts ^22^, as well as reduced training responsiveness between individuals ^32^. Another possibility is that the timing of transcriptional responses to acute exercise in the trained state experiences a “phase shift” relative to untrained muscle, and that this is related to “priming” by altered DNA methylation. Such a phenomenon was recently described in mouse muscle with endurance exercise training ^113^. There is a need for time course studies in humans evaluating the molecular responses to RE in the trained versus untrained state so that correct interpretations can be made regarding differential expression of genes (such as *MYC*) versus differential timing. Regardless, our current findings reinforce the evidence for acute exercise responses operating under the control of early methylation events ^68,114^ and support data suggesting that methylation changes are central to exercise training adaptations in humans ^7,110,115,116^.

Our work points to MYC as a key player in controlling hypertrophic adaptation to exercise in skeletal muscle. In mice, *Myc* is actively transcribed and enriched in myonuclei during mechanical overload ^24^. A single pulse of *Myc* in skeletal muscle markedly alters the transcriptome (soleus>plantaris>quadriceps) ^24,27^ and shifts the DNA methylation landscape ^27^. In humans, MYC may also act directly on rDNA after RE to influence ribosome biogenesis ^25^, consistent with MYC’s occupation of the rDNA promoter during load-induced muscle hypertrophy in mice ^100^. MYC controls ribosome biogenesis as well as skeletal muscle protein synthesis independent from mTORC1 activation ^39,117^, but its ability to drive muscle growth has been unclear ^43,117^. It is likely that chronic induction of MYC in muscle is detrimental to mass and function, similar to what occurs with prolonged chronic mTORC1 activation ^45,46^. By using a doxycycline-inducible and pulsatile model of MYC induction in skeletal muscle, we show for the first time that MYC can promote growth of skeletal muscle mass in the soleus of female mice. Our prior and current work suggests this hypertrophy could be due to ribosomal regulation – biogenesis and/or specialization ^24,27^. This larger muscle size may also be attributable to several other processes such as actin folding and autophagy. Transcriptome data from MYC induction in the soleus muscle indicates that a variety of other processes contribute to hypertrophy since >1300 genes are altered by a single MYC pulse ^27^. Our findings will lead to further detailed investigation on how MYC supports long-term muscle anabolism at the molecular, signaling, and cellular level across muscle types, fiber types, sexes, and age.

Collectively, the time course of -omics responses to RE in healthy untrained humans, alongside the repeated biopsy control group, is a valuable resource to the skeletal muscle field. Our results define the molecular landscape after exercise at high temporal resolution and will help inform the design of future human exercise studies. We detail the interplay between the methylome and transcriptome, identify MYC as a key component of the RE response, and show that MYC is sufficient for muscle growth. This study complements previous and ongoing efforts at defining the acute muscle -omics responses to RE in humans ^26,65,118^ to uncover novel molecular regulators of hypertrophic physical activity. We lay the groundwork for future investigations that will further expand on how transcriptional regulators such as *MYC* control muscle mass and adaptation in skeletal muscle.

## Methods

### Ethical approval

The regional Ethical Review Board in Linköping (2017/183-31) approved the study protocol for the human intervention. The volunteers received oral and written information about the study, and subsequently provided their informed consent prior to study enrollment. The study protocol conformed with the Declaration of Helsinki. IACUCs at the University of Arkansas (UA, AUP 21038) approved all animal procedures. Mice were housed in a temperature and humidity-controlled room, maintained on a 12:12 hour light:dark cycles, and food and water were provided *ad libitum* throughout experimentation. All animals were sacrificed via cervical dislocation under deep anesthesia with inhaled isoflurane.

### Volunteers

A subset of thirteen volunteers was chosen for analysis from previously published studies ^25,119^. Eight recreationally active volunteers were analyzed from the RE group, and five from the CON group. The volunteers in the RE group had a mean age of 32 ± 5 years, height of 181 ± 9 cm, weight of 83 ± 8 kg, and body mass index (BMI) of 25.3 ± 2.0. The corresponding values for the CON group were an age of 30 ± 4 years, height of 177 ± 5 cm, weight of 85 ± 12 kg, and a BMI of 27.3 ± 3.6.

### Experimental protocol

The experimental protocol has been described elsewhere ^25,119^. In short, volunteers were instructed to not partake in any strenuous physical activity for the legs for three days prior to the intervention. Following an overnight fast, subjects consumed a breakfast consisting of a standardized amount of liquid formula supplying 1.05 / 0.28 / 0.25 grams of carbohydrates / protein / fat per kg of body weight (Resource Komplett Näring, Nestlé Health Science, Stockholm, Sweden). Skeletal muscle biopsies were collected from the vastus lateralis, using a Bergström needle with manually applied suction ^120^. Ninety minutes after breakfast, volunteers started a 45-minute standardized RE session. The RE session consisted of a short warm-up on a cycle ergometer, followed by four sets at 7RM load with two min of rest using both leg press and leg extension machines. Muscle biopsies were collected one hour after breakfast (Pre), as well as 30 minutes and three hours after RE completion. Immediately following the three-hour biopsy, another portion of the standardized liquid formula was administered for lunch to the volunteers (2.1 / 0.56 / 0.5 grams of carbohydrates / protein / fat per kg of body weight). Eight hours after RE ended, another muscle biopsy was collected, whereby the volunteers were sent home overnight. At home, volunteers were instructed to eat a standard dinner (a balanced meal of approximately 25% of a protein source and equal distribution of carbohydrate sources and greens) in the evening. Another administered standardized liquid formula breakfast was ingested the following morning 90 minutes prior to reporting to the laboratory for the final muscle biopsy sampling 24 hours after RE completion (breakfast ingested 2 hours prior to biopsy sampling). The experimental protocol is depicted in Figure 1A (CON) and Figure 2A (RE). The sampling timepoints for the CON group was matched to the exercise group.

### RNA extraction, sequencing, and analysis

Approximately 25 mg of muscle tissue was homogenized in TRI Reagent (Sigma-Aldrich, St Louis, MO, USA) using a Bullet Blender Tissue Homogenizer (Next Advance, Troy, NY, USA). An RNA supernatant phase was then isolated using bromochloropropane and centrifugation. Next, the RNA phase was processed using Direct-zol filter columns (Zymo Research, Irvine, CA, USA). Finally, the RNA was treated with DNAse and eluted in DEPC-treated water prior to storage at -80°C. Concentration and purity of the RNA was determined using a BioTek PowerWave XS microplate reader (BioTek Instruments Inc., Winooski, VT, USA). Library preparation of mRNA was done using Poly A enrichment, followed by RNA sequencing by an Illumina NovaSeq 6000 (150 bp paired-end sequencing; Novogene Corp. Inc., Sacramento, CA, USA).

Quality control of raw sequencing reads was performed by removing adaptors and low-quality reads. Subsequently, the reads were aligned to the human reference genome (GRCh38.p12) using HISAT2 (2.0.5)^121^, and the quantification of reads mapped to each gene was conducted using featureCounts (1.5.0-p3) ^122^.

Raw counts were used as inputs for the downstream analysis in R platform (Version: 4.1.0). After filtering out genes with low expression, DESeq2 (1.42.0) was used for the normalization and differential analyses in the comparison between different timepoint groups ^123^. Genes with a false discovery rate (Benjamini-Hochberg method) adjusted *p*-value < 0.05 were identified as differentially expressed genes (DEGs). We have used org.Hs.eg.db (3.13.0) as reference for the annotation of human genes (org.Hs.eg.db: Genome wide annotation for Human. R package version 3.8.2.) ^124^. To determine gene expression patterns among the DEGs, we computed z-score per gene along different timepoints for each group separately and employed the Euclidean hierarchical clustering method to identify clustered genes. The total number of clusters was determined empirically. Raw and processed files have been deposited in the GEO database (GSE252357).

### DNA methylation data processing and statistical analysis

We previously published reduced representation bisulfite sequencing (RRBS) for ribosomal DNA (rDNA) ^25^. Here, we used this RRBS dataset for global DNA methylation analysis^125^. Quality control and adapter sequence trimming were performed using FastQC and Cutadapt, respectively as parts of the Trim Galore wrapper. Low quality base calls (Phred score <20) were removed prior to trimming adapter sequences.

Bismark aligner was used to align the sequence reads to the bisulfite-converted GRCh38 genome prior to data processing. Coverage (.cov) files produced from Bismark aligner were used for data analysis in the methylKit R package^126^. MethylKit was used to pool samples into their respective groups to maximize read coverage across the genome using a minimum read cutoff of 10 reads per base, and minimum base coverage of 1 sample per group. Differentially methylated regions (DMRs) were determined by genomic ranges for every gene promoter as defined by the hg38.bed file obtained from NCBI. Fisher’s exact test with sliding linear model (SLIM) correction for false discovery^127^ was used to qualify both differentially methylated sites and differentially methylated promoters within the dataset. Percent methylation and percent differential methylation were then obtained from methylKit following analysis.

### Sequencing, transcriptional regulator, and BETA analysis

The up- and down-regulated DEG (adj. *p*<0.05) were initially separated. All DEGs from 30 minutes, 3-, 8-, and 24 hours post-exercise biopsies were collapsed into one DEG list across the 24-hour recovery period for up- and down-regulated genes, respectively. Gene pathway enrichment analyses were conducted on the collapsed gene lists using Enrichr (https://maayanlab.cloud/Enrichr/ 2023-Aug-16) with the 2023 gene ontology (GO) database as our cross reference ^55–58^. Analyses were run using the list of sequenced genes as background correction according to Stokes et al. ^54^. The number of genes and adjusted p-values of selected enriched pathways are presented in Figure 2. The time course analysis for each pathway is based on the proportion of DEGs within each pathway in the current data set expressed at each biopsy time point relative to Pre. Thus, the number of DEGs within a specific pathway per timepoint is divided by the number of DEGs within the same pathway from the pooled 24-hour DEG list, described above.

Landscape In Silico deletion Analysis (LISA) was run according to the recommended procedures as reported by Qin et al. ^44^, consistent with our previous work ^24,27^. In brief, DEG lists (adj. *p*<0.05) were run using software on http://lisa.cistrome.org (2023-Aug-20). If the number of DEGs was above 500, the analysis was run locally using the command line. Lisa analysis was performed on the collapsed upregulated 24-hour gene list, and on up-regulated genes expressed at different timepoints. The Cauchy combination *p*-value test was used to determine the overall influence of MYC. Overlapping DEGs between the human RE response and the MYC mouse was analyzed using Venny2.1 (https://bioinfogp.cnb.csic.es/tools/venny/index.html 2023-Aug-23).

We presented the incorporation of RRBS and RNA-seq data using BETA in a previous publication ^67^. BETA is a software that provides an integrated analysis of transcription factor binding to genomic DNA and transcript abundance using chromatin immunoprecipitation sequencing (ChIP-seq) and transcriptomics (RNA-seq) datasets^128^. BETA takes into consideration the distance of the regulatory element relative to the transcription start site (TSS) by modeling the effect of regulation using a natural log function termed the regulatory potential (Eq. 1), as described previously by Tang and colleagues^129^. CpG islands were converted into “methylation peaks” similar to transcription factor binding peaks, which is built using the GRCm39 CpG island bedfile downloaded from the UCSC genome browser. Only genes differentially expressed with adjusted *p* < 0.05 from RNA sequencing analysis were included as input for gene expression. The BETA basic command was run with the following parameters “-c 0.05 --df 0.05 --da 500”.

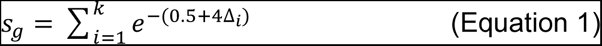

In the current study, a CpG island (κ) within 100 kb of TSS of gene (*g*) is included in the calculation for the regulatory potential score (*s*). The distance between the CpG island and the TSS is expressed as a ratio relative to 100 kb (Δ). The scoring is weighted based on the distance from the TSS (higher for smaller distances, lower for larger distances).

### BETA integration pathway analysis

Pathway analysis was performed using the enrichR R package. Up- and down-target genes resulting from BETA integration of DNA methylation and RNA-sequencing datasets were included in independent up and down gene sets. The 2023 Gene Ontology database Biological Processes was used to annotate up and down gene sets and determine enriched ontologies for each gene set. GOplot R package was used to combine log2 fold change for each gene with their respective enriched pathways.

### Generation of HSA-MYC mice and in vivo pulsatile overexpression experiments

Human skeletal actin reverse tetracycline transactivator tetracycline response element “tet-on” MYC (HSA- MYC) mice were generated as previously described ^24,27^. A subset of mice (n=6) was crossed with tet-on green fluorescent protein mice for myonuclear isolation experiments not presented here. For all MYC experiments, littermate mice (HSA or HSA-GFP) were controls; all mice were heterozygous for each transgene. At four months of age, control (n=7) and MYC overexpressing female mice (n=9) were given doxycycline in drinking water with sucrose (0.5 mg mL^−1^ with 2% sucrose) for 48 hours. All mice were then given un-supplemented drinking water for the remaining 5 days of the week. This dosing strategy was repeated 5 total times. All mice were euthanized 24 hours following the final doxycycline treatment. Some mice were used for analyses not described here, so the histology results are from n=4 control and n=6 MYC induction mice. Mice were euthanized in the morning (before 10:00 AM) and all tissues were harvested, weighed, and frozen in liquid nitrogen-cooled isopentane using optimal cutting temperature compound. The average mass of both muscles for every mouse is presented.

### Immunohistochemistry

Fiber cross sectional area and fiber type analyses on the soleus muscles were performed as previously described ^76,103^. Briefly, 8 µm sections were cut using a cryostat and air dried for ≥1 hour. Primary antibodies for dystrophin (1:100, ab15277, Abcam, St. Louis, MO, USA) and MyHC 1 (BA-D5, Developmental Studies Hybridoma Bank, Iowa City, IA, USA) were applied for ≥4 hours in a PBS cocktail. After several PBS washes, isotype-specific secondary antibodies were applied for 1 hour. Following several PBS washes, 4′,6-diamidino- 2-phenylindole (DAPI) was applied, and the slides were mounted with cover slips using a 50/50 solution of PBS and glycerol. Muscle cross sections were imaged using a Zeiss AxioImager M2. Fiber cross sectional area, fiber number, and fiber type distribution was analyzed using MyoVision ^130,131^, as we have previously described.

### Statistical considerations

Figures were generated using GraphPad Prism version 7.00 for Mac OS X (GraphPad Software, La Jolla, CA), Rstudio, BioRender, and Affinity Designer 2.3. Data presented as mean ± standard deviation of mean unless otherwise stated. Average muscle weights for each animal (mean of both muscles) and histology data were analyzed using two-tailed dependent t-tests with *p*<0.05. For all -omics analyses, Benjamini-Hochberg adjusted p values (adj. *p*<0.05) were utilized.

## Acknowledgments

Thank you to the volunteers for their willingness to participate in this research and contribute their tissue and time. Thank you to Kate Mamiseishvili, PhD, Dean of the College of Education at the University of Arkansas for resources that enabled the rapid completion of the histology. Thank you to Charlotte Peterson, PhD, and John McCarthy, PhD, of the University of Kentucky for their support of this work.

## Authors’ Contributions

KAM and FVW conceptualized the study.

SE, RGJ, NTT, PJK, SK, CSP, FMC, LNS, PRJ, LZ, VCF, YW and KAM generated and analyzed data.

SE, RGJ, KAM, and FVW interpreted the results.

SE and KAM wrote the manuscript with assistance from RGJ and input from FVW.

SE, RGJ, PJK, and PRJ generated figures.

RFG, JN, BA, and FVW oversaw and performed human exercise intervention.

RGJ, SK, and KAM performed mouse experiments.

SE, NPG, CSF, JTL, BA, KAM, and FVW provided resources and oversight.

All authors provided feedback and final approval of the manuscript.

## Data Availability

RNA-sequencing data were deposited in GEO under accession GSE252357. RRBS was deposited in association with our previous publication (Figueiredo et al. 2021) ^25^.

---

**Supplemental Figure 1.**
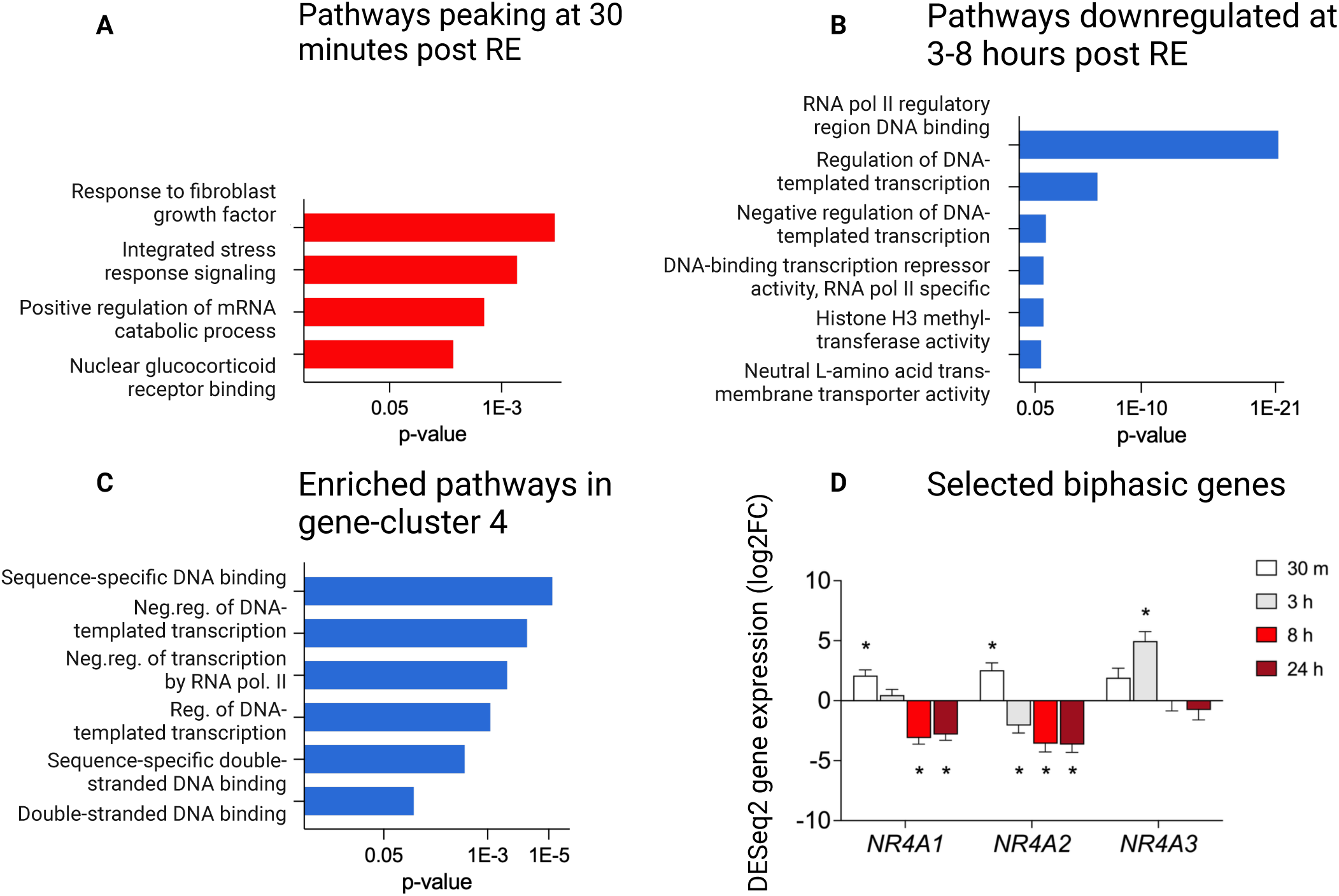
Targeted pathway enrichment analysis. Pooled gene ontology (GO) biological processes and molecular function pathways. (A) Targeted analysis of pathways peaking at 30 minutes post resistance exercise (RE). (B) Targeted analysis of pathways significantly enriched in genes downregulated 3-8 hours post RE. (C) Targeted analysis of biphasic DEGs composing cluster 4, as presented in figure 2D. (D) Gene expression of selected genes with a biphasic gene expression pattern, up early/down late. * = p<0.05 vs Pre values. Neg. = Negative, Reg. = Regulation.

**Supplemental Figure 2.**
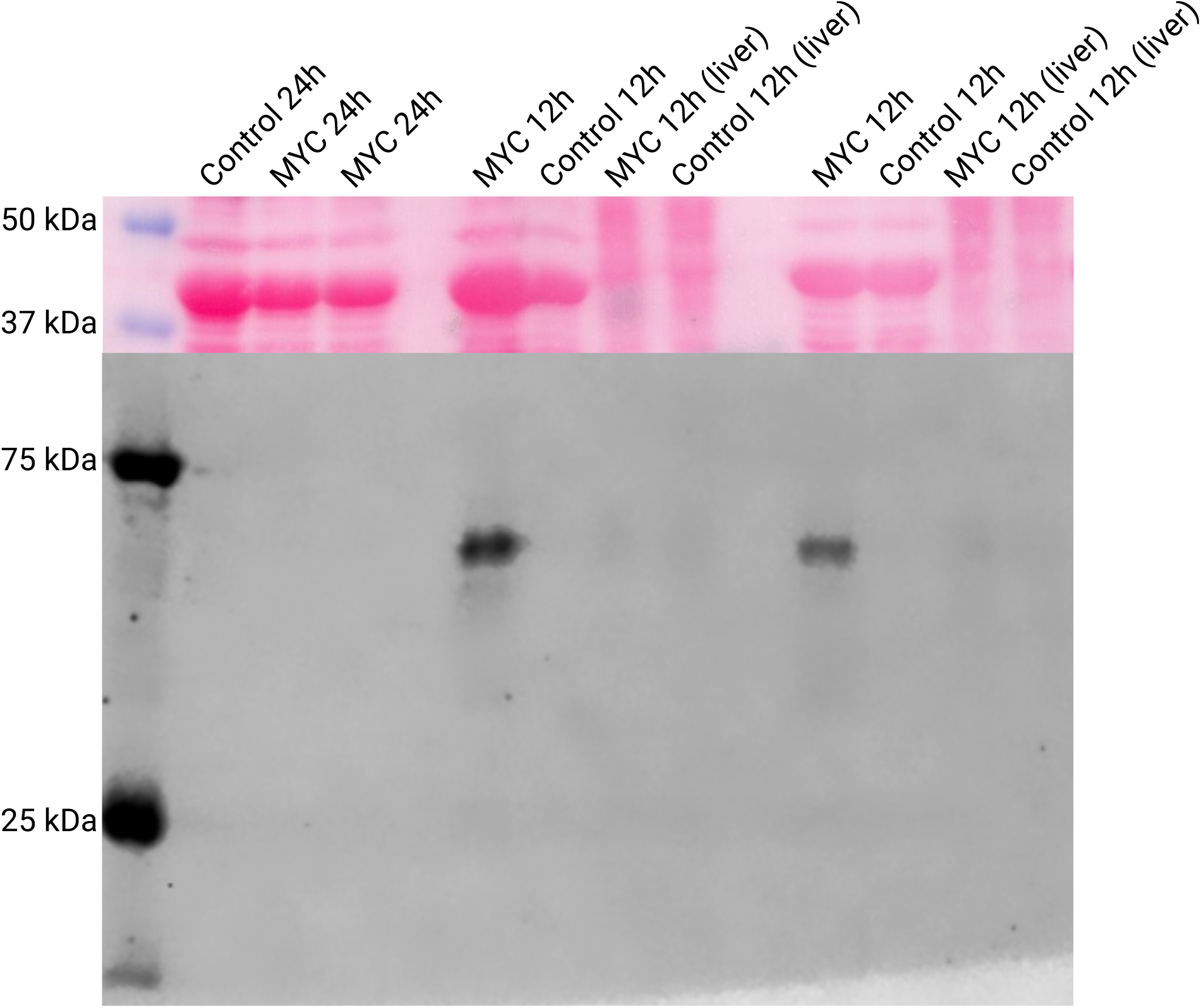
Levels of MYC protein return to baseline after 24 hours cessation supplemented water administration. HSA-MYC and HSA littermate control mice were treated with doxycycline in drinking water for 12 hours, followed by a 24-hour treatment with un-supplemented drinking water. Quadriceps muscle was harvested in the morning and western blots for MYC were carried out as described in our previous publication ^27^. Lanes 1-3 show one doxycycline-treated HSA and two HSA-MYC mice following 12 hours of doxycycline and 24 hours of un-supplemented water. MYC is not expressed in HSA mice and levels return to baseline levels by 24 hours in HSA-MYC mice (no MYC protein detectable). Lanes 4 & 8 show quadriceps muscle expression in HSA-MYC mice after 12 hours of treatment with doxycycline followed by a 12-hour chase (positive control, see Jones III et al. 2022). Lanes 5 & 9 show the HSA control mice after the same 12-hour chase (negative control). Lanes 6 & 10 and 7 & 11 show MYC expression in liver of corresponding animals (another negative control).

**Supplemental Figure 3.**
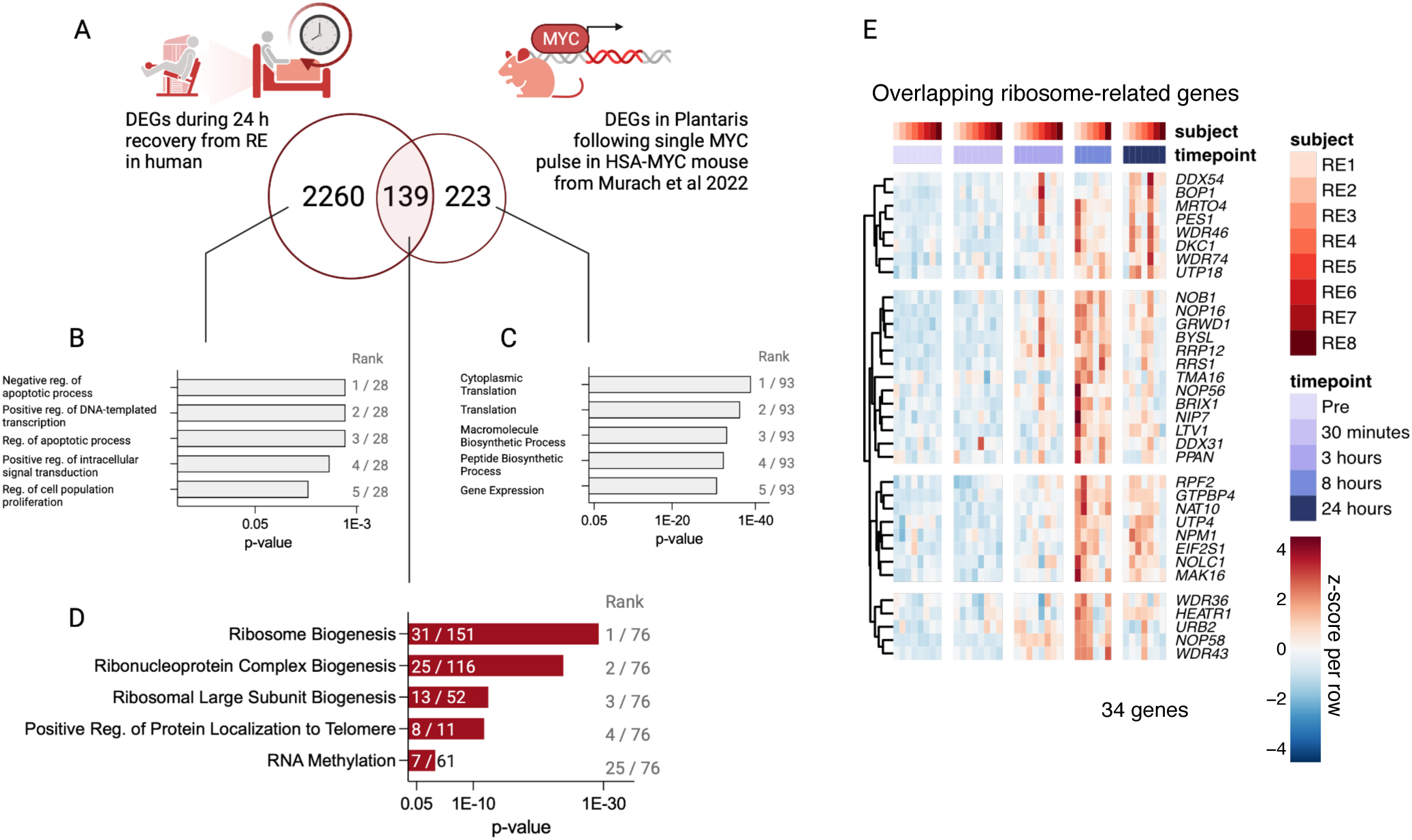
Transcriptional similarities between human RE recovery and MYC overexpression in mouse plantaris muscle. (A) Comparison of up-regulated DEGs across 24 hour of RE recovery in humans (n=8) vs plantaris muscle from MYC-overexpressing mice from Murach et al (2022). (B-D) Top pathways (GO: Biological processes) based on DEGs in B) the human exclusive gene list, C) MYC mouse exclusive gene list, and D) overlapping gene list, respectively. Pathways are ranked according to their adj. *p*- values. (G) Heatmap showing DEG pattern for ribosome-related genes overlapping human RE response to a MYC response in mouse plantaris muscle. Genes retrieved from all five pathways presented in S3D.

